# Sodium valproate protects against neuronal ferroptosis in epilepsy via suppressing lysyl oxidase

**DOI:** 10.1101/2020.05.31.126227

**Authors:** Qin Li, Qiu-Qi Li, Ji-Ning Jia, Zhao-Qian Liu, Hong-Hao Zhou, Wei-Lin Jin, Xiao-Yuan Mao

## Abstract

**Background and purpose:** Epilepsy is a chronic neurological disease that is characterized by repetitive seizures. Seizures-related complications such as cognitive deficits, anxiety and sleep disorders seriously impact the life quality of patients. Antiepileptic drugs are widely used for the treatment of epilepsy. Sodium valproate is served as the first-line antiepileptic drugs and possesses various pharmacological effects on the brain. Sodium valproate exerts neuroprotective effects in acute nervous system diseases such as ischemic brain damage by inhibiting oxidative stress. However, the mechanism of neuroprotection of sodium valproate in epilepsy is unclear. Lysyl oxidase (Lox) is a monoamine oxidase that acts on extracellular matrix collagen and elastin and it can promote accumulation of oxidative stress. Our previous studies have confirmed that Lox is involved in ferroptosis, a novel iron-dependent and lipid peroxidation-mediated cell death pathway, during epilepsy. In this study, we would like to investigate whether sodium valproate can exert neuroprotective effects on kainic acid-induced epileptic seizures by inhibiting Lox-mediated ferroptosis.

**Methods:** Epileptic mouse models were established by intracranial injection of 250 ng/μl kainic acid on right hippocampus. Sodium valproate and ferroptosis inhibitors were administrated by intraperitoneal injecting. The epileptic behavior of the mice within 4 hours was recorded after intracranial injection of kainic acid. Mouse hippocampus was acquired to analyze the mRNA expression of prostaglandin-endoperoxide synthase 2 (*PTGS2*) and the production of 4-hydroxynonenal (4-HNE). In vitro, the protective effects of sodium valproate on glutamate-induced HT22 cell damage model was assessed by PI/Hoechst staining; The levels of *PTGS2*, 4-HNE and lipid ROS were analyzed by RT-qPCR, western blot and flow cytometry, respectively. RT-qPCR and Western blot analysis the mRNA and protein expression of Lox in the glutamate-induced HT22 cell damage model. The Lox overexpression model was established by intracranial injection of AAV on right hippocampus.

**Results:** Pretreatment with sodium valproate and ferroptosis inhibitors could significantly alleviate the epileptic seizures in the kainic acid induced epilepsy mouse model. Western blot and RT-qPCR results showed that sodium valproate and ferroptosis inhibitors significantly inhibited the levels of 4-HNE and *PTGS2*. PI/Hoechst staining showed that 1 mM sodium valproate exerted protective effect on glutamate-induced HT22 cell injury model. There was no significant difference observed between sodium valproate and ferroptosis inhibitors co-intervention group and sodium valproate intervention group on glutamate-induced cell injury model. And sodium valproate could significantly inhibit the production of lipid reactive oxygen species and 4-HNE. The expression of Lox was significantly increased in the glutamate-induced HT22 cell injury model, which could be reversed by pretreatment of sodium valproate. And β-aminopropionitrile (a specific inhibitor of Lox) could inhibit ferroptosis induced by glutamate, as well as ameliorate the epileptic seizures in the kainic acid induced epilepsy mouse model. Pretreatment with sodium valproate could not ameliorate the epileptic behavior in the Lox-overexpression mice. Western blot analysis showed that sodium valproate could not suppress the production of 4-HNE in kainic acid induced epileptic mice model.

**Conclusions:** The neuroprotective effect of sodium valproate in epileptic seizures is closely related to the inhibition of ferroptosis. The inhibition of ferroptosis is involved in the neuroprotective effect of sodium valproate on glutamate-induced HT22 cell damage model. Sodium valproate may exert neuroprotective effects in kainic acid-induced epileptic seizures by abrogating Lox-mediated ferroptosis.

## 1. Introduction

Epilepsy is a chronic neurological disease characterized by spontaneous recurrent seizures^[1]^. It is considered to be the second most common disease after stroke^[2]^. According to statistics, about 70 million people around the world are shrouded in the shadow of epilepsy^[3]^. Several syndromes including cognitive decline and depression, usually occur in the epileptogenesis^[4,5]^, which have a detrimental effect on the life quality of patients. The prevalence of major depression in the general population is 4.9%~7% while the prevalence of depression in patients with epilepsy is as high as 11%−60%^[6]^. The treatments for epilepsy involve medical treatment, surgical treatment, diet therapy and hormonal interventions, among of which medical treatment is offered as the primary treatment^[6,7]^. Valproic acid (VPA), a traditional broad-spectrum antiepileptic drug, is known to have diverse pharmacological effects, such as prolongation of sodium-channel inactivation^[8]^, elevation of GABA content in the brain^[9]^, inhibition of HDAC activity^[10]^, reduction of reactive oxygen species (ROS) production^[11]^. However, its mechanism for the antiepileptic action remains poorly understood.

Recently, oxidative stress has been shown to be related to the mechanism for the neuroprotective of VPA. In a traumatic brain injury model, VPA increases antioxidative factors through activation of the Nrf2/ARE signaling pathway ultimately leading to neuroprotection ^[12]^. Oxidative stress is a situation that introduces a high production of oxidants or a low level of antioxidants, which results in an imbalance between oxidant and antioxidant systems causing free radical damage. Accumulating evidence indicates that oxidative stress is critical to the etiology of many diseases, especially neurodegenerative diseases and cancers^[13]^. Carbonyl protein, a potential oxidative damage biomarker, are highly accumulated in the Alzheimer’s disease brain and are localized to paired helical filaments and amyloid plaques, hallmarks of Alzheimer’s disease, implying that these protein modifications may play a causal role in the progression of the neurodegenerative disease^[14]^. Besides, oxidative damage of lipids is crucial in the development of neurodegenerative diseases due to the abundant polyunsaturated fatty acids in the lipid bilayer of the brain^[15]^. And lipid peroxidation can lead to cellular apoptosis or programmed cell death in the nervous system^[16]^. So, it can be speculated that lipid peroxidation-mediated oxidative damage may be an important type of injury in neuronal damage. In addition, it was reported that VPA could protect neuron cells from damage through inhibiting the lipid peroxidation and protein oxidation^[17]^. Meanwhile, with the continuous study of the forms of cell death, more and more studies have shown that lipid peroxidation can involve the progress of ferroptosis^[18]^. This suggests that VPA may produce a neuroprotective effect by inhibiting ferroptosis regulated by lipid peroxidation. Ferroptosis is a new way of cell death that is different from apoptosis, necrosis and autophagy. Ferroptosis is an iron-dependent form of programmed cell death. Its main characteristic are the accumulation of lipid peroxides, smaller mitochondria, and increased bilayer density^[19]^. Similar to apoptosis and autophagy, ferroptosis is also associated with a variety of diseases. Current studies have found that the pathological cell death of neuroprogressive diseases (such as Alzheimer’s, Huntington’s, and Parkinson’s diseases), stroke, ischemia-reperfusion injury, traumatic brain injury, renal injury, and iron metabolism-related diseases are all closely associate with ferroptosis^[20–23]^. Ferroptosis inhibitor ferrostatin-1 can protect cell death in cellular models of Huntington’s disease^[24]^. Recently, Do Van B etc. confirmed that ferroptosis was a new way of cell death in Parkinson’s disease^[25]^. Ferroptosis also makes contribution to the neuronal loss in Alzheimer’s disease^[26]^. More and more studies have shown that ferroptosis is involved in neurological diseases. The mechanisms linking ferroptosis to neuron cell death in epilepsy, however, has not yet been clarified^[27]^. In summary, it is reported that VPA can achieve a neuroprotective effect by inhibiting lipid peroxidation, and lipid peroxidation can mediate the occurrence of ferroptosis. And whether ferroptosis is involved in the antiepileptic effect of VPA is still unknown. More evidence is needed to elucidate the role of ferroptosis in the antiepileptic effects of VPA. In this study, our main purpose is to investigate whether ferroptosis, a new form of cell death, may be used as a potential mechanism for the antiepileptic effect of VPA, thus providing a new therapeutic strategy for the treatment of epilepsy.

## 2. Materials and methods

### 2.1 Cell culture and drug treatments

Immortalized mouse hippocampal HT22 cells were grown in DMEM containing 10 % (v/v) FBS and 1% penicillin-streptomycin at 37 °C in a 5 % CO_2_ atmosphere. After that, hippocampal HT22 cells were pretreatment with 12.5 μM Ferrostatin-1 (Fer-1) (S7243, Selleck, USA), 50 μM deferoxamine mesylate (DFO) (D9533, Sigma, USA) or 1 μM liproxstatin-1 (Lipo-1) (S7699, Selleck, USA) for 2 h. Then 5 mM glutamate (G8415, Sigma, USA) or 500 nM erastin (S7242, Selleck, USA) were added into medium for 8 h.

### 2.2 Microscopic analysis for cell viability by PI and Hoechst 33342 double staining

Briefly, HT22 cells were seeded into the 24-well plates with a proper density. Then, cell death caused by drugs were detected by double staining the cells with 5 μl Hoechst 33342 (23491-52-3, Biotech, China) and Propidium iodide (PI) (P4170, Sigma, USA) for 15 min at 37 °C. Images of the staining were captured with a fluorescent microscope (Olympus, Tokyo, Japan). The total number of PI positive cells in four different representative fields per well were quantified for each treatment group to estimate the percent of PI positive cells out of the total cell number.

### 2.3 Quantitative Real-time Polymerase chain reaction

We used Trizol (15596-018, Invitrogen, USA) to extract total RNA following the provider’s protocol. Then the synthesis of cDNA was performed with Prime Script RT reagent Kit (RR047A, TaKaKa, China). RT-PCR was taken using a Light Cycler 480 system (Roche, Germany) with double-stranded DNA dye SYBR Green (RR091A, Takara, Japan). Complementary DNA from various cell samples or hippocampal tissue samples were amplified with specific primers (Lox, Forward: 5′-TCTTCTGCTGCGTGACAACC-3′, Reverse: 5′-GAGAAACCAGCTTGGAACC AG −3′; PTGS2, Forward: 5’-GGGAGTCTGGAACATTGTGAA-3’, Reverse: 5’-GTGCACATTGTAAGTAGGTGGACT-3’; Actin, Forward: 5′-GTGACGTTGAC ATCCGTAAAGA-3′, Reverse: 5′-GCCGGACTC ATCGTACTCC-3′). The cycling process of PCR real time were programmed as following: pre-incubation step (95 °C, 30 s × 1 cycles), amplification step (denaturation 95 °C, 5 s; annealing 55 °C, 30 s; and extension at 72 °C, 30 s × 40 cycles), followed by a melting curve analysis (95 °C, 0 s; 65 °C, 15 s; and fluorescence acquisition at 95 °C, 0 s). The comparative threshold cycle method (2^−ΔΔCt^) was used to quantify the mRNA expression of genes.

### 2.4 Western blot

Mouse hippocampus or cells samples were lysed using radio immunoprecipitation assay (RIPA) lysis buffer as previous instructions. Briefly, grinding the mouse hippocampus with a mortar and lysed in ice-cold RIPA lysis buffer with protease inhibitor, while cells were directly lysed in ice-cold RIPA lysis buffer. After 5 s sonication for three times, protein concentration was determined using the bicinchoninic acid (BCA) protein assay reagent (P0006, Beyotime Biotechnology, China). The protein samples (20 μg) were separated by sodium dodecyl sulfate-polyacrylamide gel electrophoresis and transferred onto polyvinylidene fluorid membranes. After blocking in tris buffered saline tween (TBST) buffer with 5% non-fat milk for 1 h, the membranes were then incubated with primary antibodies (Lox, rabbit, 36 KDa, ab31238, 1:600 dilution, Abcam; 4-hydroxynonenal (4-HNE), rabbit, 17-76 KDa, ab46545, 1:3000 dilution, Abcam; β-actin, mouse, 43 KDa, A5441, 1:10000 dilution, Sigma) at 4 °C overnight. After being washed in TBST buffer for three times, the membranes were further incubated with secondary antibodies (mouse, 1:10000 dilution, A9044, Sigma; rabbit, 1:10000 dilution, A9169, Sigma). Immunoreactive bands were visualized by ChemiDoc XRS+ imaging system. β-actin was used as an internal control. For the quantification of the protein bands, Image J software (National Institutes of Health, USA) was used to obtain the densitometric values.

### 2.5 Measurement of malondialdehyde (MDA) levels

The MDA level was determined by using the Lipid Peroxidation MDA Assay Kit (S0131, Beyotime, China) according to the manufacturer’s protocol. Cells were lysed with lysis solution and centrifuged at 1600 g for 10 min to collect the supernatant, protein quantified by BCA kit. Preparation of MDA test working solution, 200 μl of working solution and 100 μl of supernatant were mixed uniformly and then heated at 100 °C for 15 min, after the end of the heating, placed in a water bath until cooled to room temperature, centrifuged at 1000 g for 10 min. In a 96-well plate, the absorbance was measured at a wavelength of 532 nm with a microplate reader and the MDA concentration of the sample was obtained according to a standard curve.

### 2.6 Analysis of lipid peroxidation

According to previous reports, BODIPY 581/591C11 (Thermo fisher, D3861) was used to measure lipid peroxidation. Briefly, cells were collected by centrifugation at 3000 g for 5 minutes after digestion from culture dish, and 500 μl of BODIPY (dilution with water to 10mM) was added to re-suspend cells. The cell suspension was vortexed for 15 seconds and then incubated at 37 °C for 15 min. In order to remove the excess BODIPY, cell suspension was centrifuged at 3000 g for 5 minutes and then re-suspended with 1000 μl HBSS. 300 μl of PBS was added to re-suspend cell, and examined by flow cytometry, the results were analyzed by Cell Quest program (BD Biosciences).

### 2.7 Intrahippocampal Kainic acid (KA) injection

All adult male C57/BL6 mice weighing 20-24 g were obtained from the Experimental Animal Center of Central South University, China, and were randomly divided into a normal control group and an experimental group. All animals were housed in cages with a 12/12 h light/dark cycle and other standard laboratory conditions, including a room temperature of 23 ± 1 °C and access to food and water ad libitum. KA-induced epileptic seizure model was performed as previously described. Briefly, a 5 μl microsyringe was filled with 1 μl solution of KA (K0250, Sigma-Aldrich, USA), and positioned in the right dorsal hippocampus (anteroposterior [AP] = −2 mm, mediolateral [ML] = −1.8 mm, dorsoventral [DV] = −2.3 mm). Mice were injected (over 2 minute) with 1 μL KA solution (250 ng), using a microsyringe. After injection, the needle was left in place for an additional 5 minutes to avoid reflux. Sham mice were injected with 1 μL 0.9% sterile NaCl. After surgery, the animals were kept under observation for 6 hours to assess the behavioral changes corresponding to seizures.

### 2.8 Statistics

Data are presented as mean ± SEM. Statistical analyses were performed using GraphPad Prism (GraphPad Prism Software, San Diego, CA, USA). For comparing 2 groups, either unpaired or paired t tests were used with Welch’s correction. For experiments with more than 3 groups, Bonferroni’s multiple comparisons test were used. If variances were different among samples, then Kruskal-Wallis test, followed by Dunn’s multiple comparisons test, was used. When same cells received multiple treatments, then repeated-measures one-way ANOVA, followed by Tukey’s multiple comparisons test, was used. P values of less than 0.05 were considered significant.

## 3. Results

### 3.1 VPA and ferroptosis inhibitors alleviate KA-induced epileptic seizures

Epileptic seizure model in mouse was established by stereotactic injection of KA into right hippocampus. Seizure score was evaluated within 4 h after injection by Racine. Mice were randomly divided into four groups as follows: control group, 100 ng/μl KA group, 250 ng/μl KA group and 500 ng/μl KA group. The experimental groups were received corresponding doses of KA, and control group was recived equal volume PBS. The behavioral changes of mice were recorded after injection. It was obvious that treatment with 100 ng/μl KA showed rhythmic nodding. Mice with the exhibition of grade 4-5 seizures were considered successful. It was found that an administration of a dose of 250 ng/μl KA and 500 ng/μl KA both resulted in obvious convulsive phenotype. However, there is high death rate in mice after injection of 500 ng/μl KA. Therefore, epileptic seizure model was established by injection of 250 ng/μl KA (Fig. 1A, Fig. 1B and Fig. 1C). Furthermore, pretreatment with VPA and ferroptosis inhibitors (DFO, Lipo-1 and Fer-1) significantly protected brain from epileptic seizures in mice as shown in Fig. 1D, Fig. 1E and Fig. 1F.

**Figure 1.**
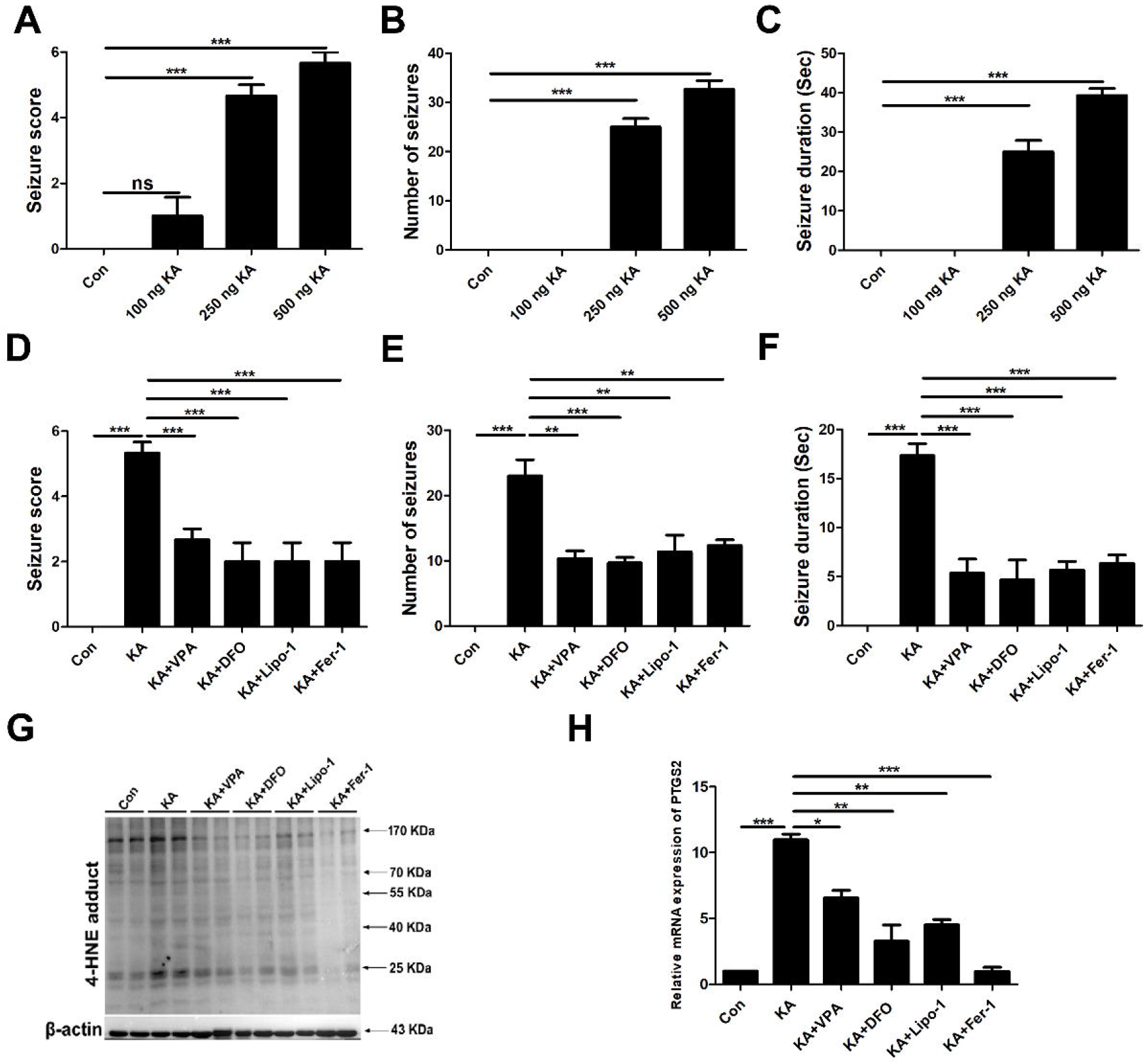
VPA and ferroptosis inhibitors significantly improved the behavioral seizures induced by kainic acid. **(A-C)** Behavioral seizures induced by different concentrations of kainic acid (100 ng, 250 ng, 500 ng) were presented as seizure score, number of seizures, and average seizure duration (Sec), n=6. **(D-F)** VPA and ferroptosis inhibitors (DFO: 100 mg/Kg; Lipo-1: 10 mg/Kg; Fer-1: 3 mg/Kg) could reduce the seizure score, number of seizures, and average seizure duration (Sec) caused by 250 ng kainic acid, n=6. **(G)** VPA and ferroptosis inhibitors (DFO: 100 mg/Kg; Lipo-1: 10 mg/Kg; Fer-1: 3 mg/Kg) could reduce the production of 4-HNE in the hippocampus tissue from kainic acid-induced epilepsy model mice, n=6. **(H)** The relative mRNA expression of *PTGS2* was reduced after VPA and ferroptosis inhibitors pretreatment, * p < 0.05, ** p < 0.01, and *** p < 0.001.

To investigate whether ferroptosis was involved in KA-induced epileptic seizures, and whether VPA exerted neuroprotective effects on KA-induced epileptic seizures by inhibiting ferroptosis, we detected mRNA expression of prostaglandin-endoperoxide synthase 2 (PTGS2) and the level of 4-hydroxynonenal (4-HNE). It was found that production of 4-HNE (Fig. 1G) and mRNA expression level of PTGS2 in KA group were significantly increased in comparison with control group (Fig. 1H). In addition, pretreatment with VPA and ferroptosis inhibitors evidently suppressed PTGS2 mRNA and 4-HNE level. Taken together, it indicates that ferroptosis was involved in epilepsy, and the protective effect of VPA in a mouse model of KA-induced epileptic seizures was related to ferroptosis inhibition.

### 3.2 Ferroptosis is involved in the protective effect of VPA on glutamate-induced HT22 toxicity

Previous studies have shown that VPA protects brain from epileptic seizures possibly via inhibition of ferroptosis. Our present work sought to explore the potential molecular mechanism by which VPA alleviated epileptic seizures. Glutamate is a kind of excitatory neurotransmitter which is widely distributed in the nervous system. Previous studies supported the notion that glutamate served as a contributor of seizures^[28]^. In addition, it was reported that high levels of glutamate in extracellular astrocytes aggravated epileptic seizures^[29]^, which also indicates that glutamate-induced toxicity could reflect the pathological process of seizures to some extent. In this study, glutamate-induced oxidative toxicity was induced in HT22 cells as this neuronal cell line is featured by deficient NMDA receptor^[30]^. In addition, studies have proved the existence of ferroptosis in glutamate-induced HT22 cell damage^[31]^. Our previous work has also confirmed that ferroptosis is the major type of cell death in glutamate-induced HT22 cell damage. Thus, the glutamate-induced HT22 oxidative toxicity model was selected to investigate the molecular mechanism underlying the neuroprotection of VPA against ferroptosis. And our prior work also demonstrates that glutamate at 5 mM for 8 h triggers significant damage in HT22 cells. In our subsequent experiments, we used 5 mM glutamate to treat HT22 cells for 8 h to construct the cell damage model. In order to determine the optimal concentration of VPA against glutamate-induced HT22 cell damage model, different concentrations of VPA (0.25 mM, 0.5 mM, 1 mM) were used to treat the cells at different times (pretreatment for 8 h, 5 h, 2 h or post for 2 h). PI/Hoechst staining showed that pretreatment with 1 mM VPA for 5 h had a significant protective effect on glutamate-induced HT22 cell damage (Fig. 2A - Fig. 2D).

**Figure 2.**
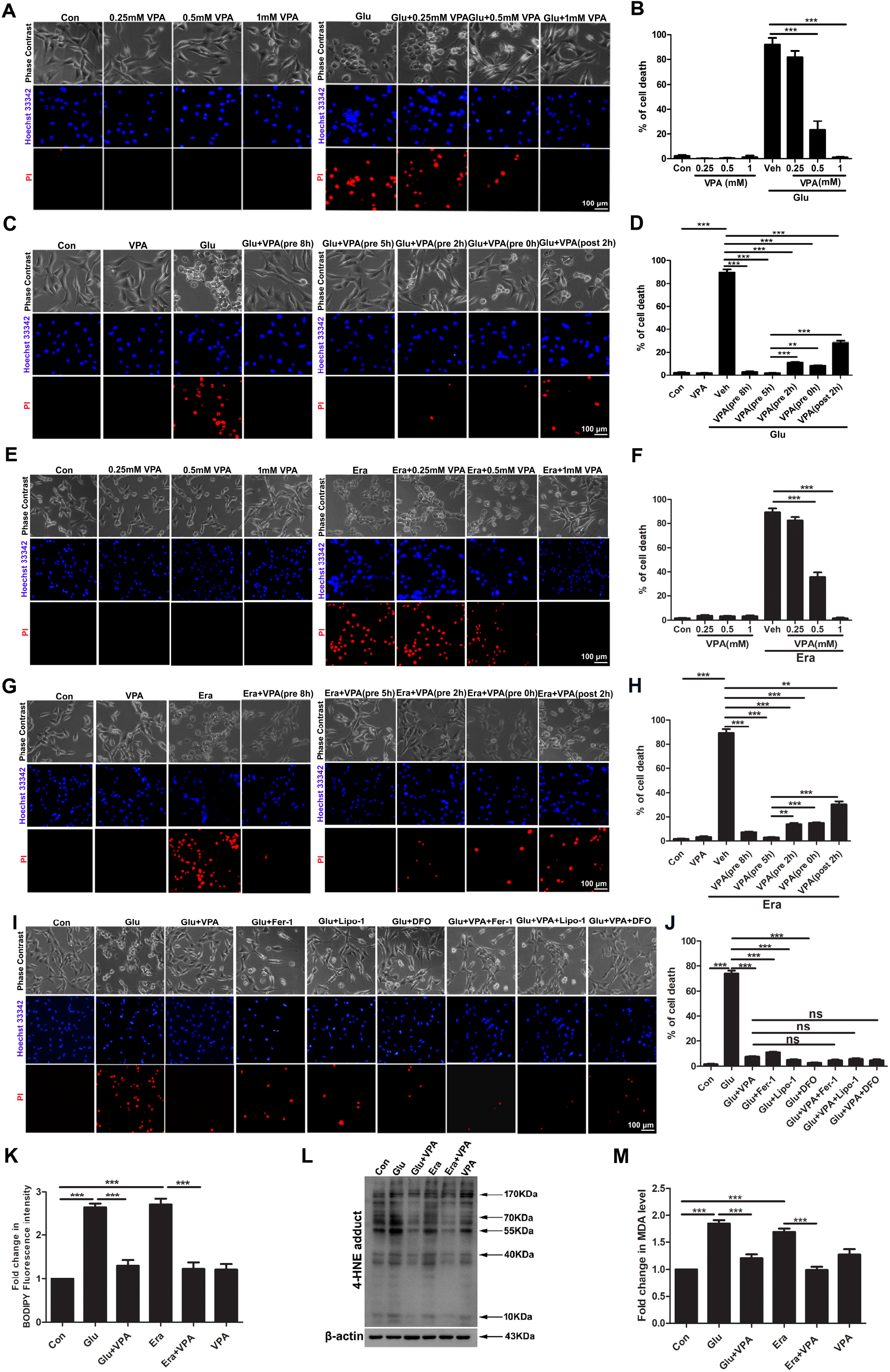
VPA prevented glutamate-induced cell death by inhibiting ferroptosis. **(A-H)** Representative images of PI/Hoechst staining showed that different concentrations of VPA (0.25 mM, 0.5 mM, 1 mM) protected HT22 cells from glutamate or erastin-induced cell death at different times (pretreatment for 8 h, 5 h, 2 h or post for 2 h), scar bar=100 μm. **(I-J)** Pretreatment with VPA (1 mM) and ferroptosis inhibitors (Fer-1: 12.5 μM; Lipo-1: 1μM; DFO: 50 μM) for 2 h could alleviate glutamate-induced HT22 cell death, scar bar=100 μm. **(K)** Pretreatment with VPA (1 mM) for 5 h could inhibit the increasing of lipid ROS in HT22 cells treated with glutamate (5 mM) or erastin (500 nM). **(L)** Pretreatment with VPA (1 mM) could reduce the production of 4-HNE in HT22 cells treated with glutamate (5 mM) or erastin (500 nM). **(M)** VPA (1 mM) could the production of MDA in HT22 cells treated with glutamate (5 mM) or erastin (500 nM), *p < 0.05, **p < 0.01 and ***p < 0.0001, n=3.

To further explore whether the protective effect of VPA on glutamate-induced HT22 cell damage is related to the inhibition of ferroptosis, three commonly used specific inhibitors of ferroptosis were selected in this study: Fer-1, lipo-1 and DFO as positive controls. Our experimental results showed that pretreatment with Fer-1 (12.5 μM), Lipo-1 (1 μM) and DFO (50 μM) for 2 h had a significant neuroprotective effect on glutamine-induced HT22 toxicity. Moreover, the protective effect of VPA and ferroptosis inhibitors co-intervention group on glutamate-induced cell damage was not significantly increased compared with VPA alone (Fig. 2I and Fig. 2J). It suggests that the protective effect of VPA on the glutamate-induced HT22 cell damage model may be related to the inhibition of ferroptosis. In order to further confirm the relationship between the protective effect of VPA on glutamate-induced HT22 cell damage model and ferroptosis, we detected the ferroptosis-related indicators in HT22 neurons following glutamate exposure and intervention with VPA. It showed elevation of lipid ROS level in glutamate-induced model group and VPA reduced the production of lipid ROS (Fig. 2K). Western blot analysis also indicated that the lipid peroxidation by-product 4-HNE was observed to be elevated in HT22 cells treated with glutamate and VPA pretreatment reversed this index (Fig. 2L). Similar results were also found in grounds of MDA level in HT22 cells following glutamate and VPA (Fig. 2M).

In order to further confirm the inhibitory effect of VPA on ferroptosis, we also investigated whether VPA inhibited erastin-induced ferroptosis. Similarly, we used different concentrations of erastin to treat HT22 cells at different times to determine the optimal condition for erastin. The results showed that 500 nM erastin treatment for 8 h caused the most obvious damage in HT22 cells. In addition, different concentrations (0.25 mM, 0.5 mM, 1 mM) of VPA to this model for different periods of time (pretreatment for 8 h, 5 h, 2 h or post for 2 h) were treated for this cell line. The results showed that pretreatment with 1 mM VPA for 5 h had a significant protective effect on erastin-induced HT22 cell damage models (Fig. 2E and Fig. 2H). At the same time, we also added VPA in the erastin-induced cell damage model to detect ferroptosis indicators, and obtained consistent results with the glutamate model (Fig. 2K - Fig. 2M). In summary, the protective effect of VPA on the glutamate-induced HT22 cell damage model was associated with ferroptosis.

### 3.3 Inhibition of Lox can reverse glutamate-induced ferroptosis in HT22 cells

Our previous studies showed that Lox was significantly upregulated in hippocampus of epileptic mice, suggesting a vital target for ferroptosis in epilepsy. Thus, our study was further to investigate whether Lox-mediated ferroptosis is involved in the protective effect of VPA on glutamate-induced HT22 cell damage model. We analyzed the alteration of Lox mRNA expression level from each group by RT-qPCR. The results showed that the relative mRNA expression of Lox was significantly promoted in glutamate-treated HT22 cells, while pretreatment with VPA could reverse this phenomenon (Fig. 3A). We further verified the above results at the protein level, and the results showed that the protein expression of Lox was significantly increased in the glutamate-induced HT22 cell injury model, which could be reversed by intervention of VPA (Fig. 3B). The above results suggested that Lox may be a target of VPA in inhibiting glutamate-induced ferroptosis. To further explore whether Lox is a potential target of VPA in inhibiting ferroptosis, we chose β-amino propionitrile (a specific inhibitor of Lox) to intervene in the glutamate-induced cell injury model. PI/Hoechst staining showed that pretreatment with BAPN (15 mM) for 30 min exerted significantly protective effect on glutamate-induced HT22 cell damage model (Fig. 3C). The study further examined the effect of BAPN on ferroptosis-related biomarkers. Flow cytometry results showed that pretreatment with BAPN significantly reduced the level of lipid ROS in glutamate-treated HT22 cells (Fig. 3D). Besides, western blots analysis indicated that the intervention of BAPN could reduce the production of 4-HNE in the glutamate-treated HT22 cells (Fig. 3E). In addition, RT-qPCR results showed that the intervention of BAPN could significantly inhibit the relative mRNA expression of PTGS2 caused by glutamate (Fig. 3F). All the above results indicated that VPA suppressed glutamate-induced neuronal ferroptosis via inhibiting Lox.

**Figure 3.**
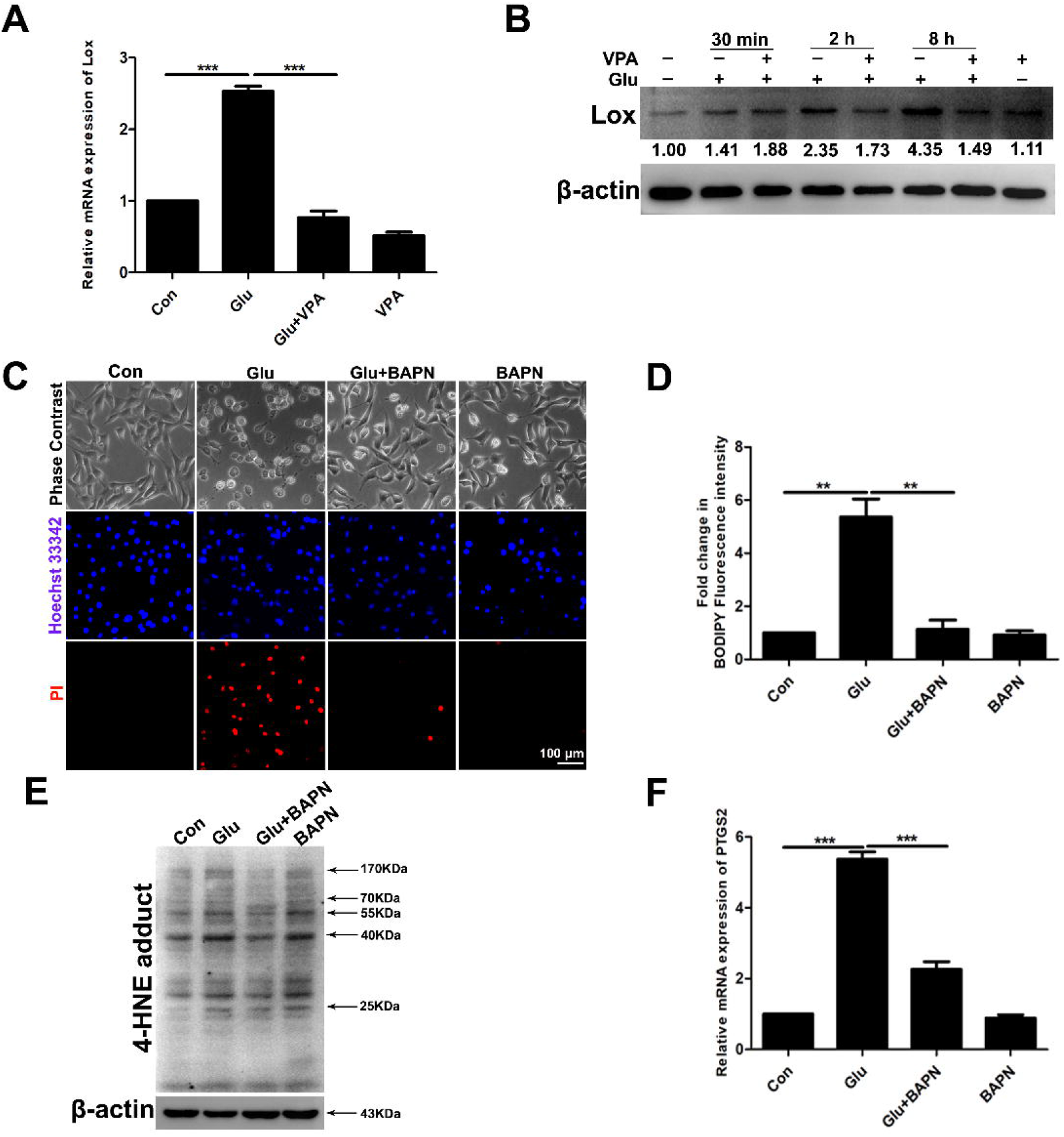
BAPN significantly inhibited glutamate-induced ferroptosis in HT22 cells. **(A)** In glutamate-treated HT22 cells, the relative mRNA expression of *Lox* was significantly reduced in VPA (1 mM) pretreatment group. **(B)** Western blot showed that pretreatment with VPA (1 mM) for 5 h significantly decreased the protein expression of Lox in glutamate treated HT22 cells. **(C)** Pretreatment with BAPN (15 mM) for 30 min could reverse cell death induced by glutamate, scar bar=100 μm. **(D)** BAPN (15 mM) significantly reduced the level of lipid ROS in glutamate treated HT22 cells. **(E)** Western blots analysis indicated that BAPN could reduce the production of 4-HNE in the glutamate treated HT22 cells. **(F)** Pretreatment with BAPN (15 mM) for 30 min could inhibit the relative mRNA expression of *PTGS2*, *p < 0.05, **p < 0.01 and ***p < 0.0001, n=3.

### 3.4 Overexpression of Lox blocks the neuroprotective of VPA in KA-induced epileptic seizures

Next, we further verified the molecular mechanism of VPA by which it inhibits ferroptosis. Similar to VPA, our present work showed that pretreatment with BAPN for two weeks significantly alleviated KA-induced epileptic seizures (Fig. 4A - Fig. 4C). It suggests that inhibition of Lox can suppress KA-induced epileptic seizures. And then, in this study, Lox overexpression virus was injected into the right hippocampus of C57 mice through stereotaxic injection to cause Lox overexpression. To verify whether Lox was successfully overexpressed, we detected the expression of eGFP in the hippocampal samples of mice by immunofluorescence (Fig. 4D). And then the protein expression level of Lox was detected by Western blot, and the results showed that the expression of Lox in hippocampus of pAAV-SYN-Lox-P2A-EGFP-3FLAG group was significantly higher than that in pAAV-SYN-EGFP-3FLAG group (Fig. 4E). It indicated that Lox overexpression C57 mice model was successfully established. Continuous intraperitoneal injection of VPA was initiated a week after viral injection, and epilepsy mouse models were established after two weeks. The results showed that VPA significantly inhibited KA-induced epileptic seizures in control group. Treatment of VPA could not reduce seizure score, number of seizures, and average seizure duration caused by KA in Lox overexpression group (Fig. 4F-H). Western blot analysis showed that VPA could significantly inhibit the production of 4-HNE caused by KA in the control group. Pretreatment with VPA could not reduce the production of 4-HNE caused by KA in Lox overexpression mice model. It indicated that Lox overexpression reversed the inhibitory effect of VPA on ferroptosis (Fig. 4I). These results indicate that VPA exerts neuroprotective effects in KA-induced epileptic seizures by inhibiting Lox-mediated ferroptosis.

**Figure 4.**
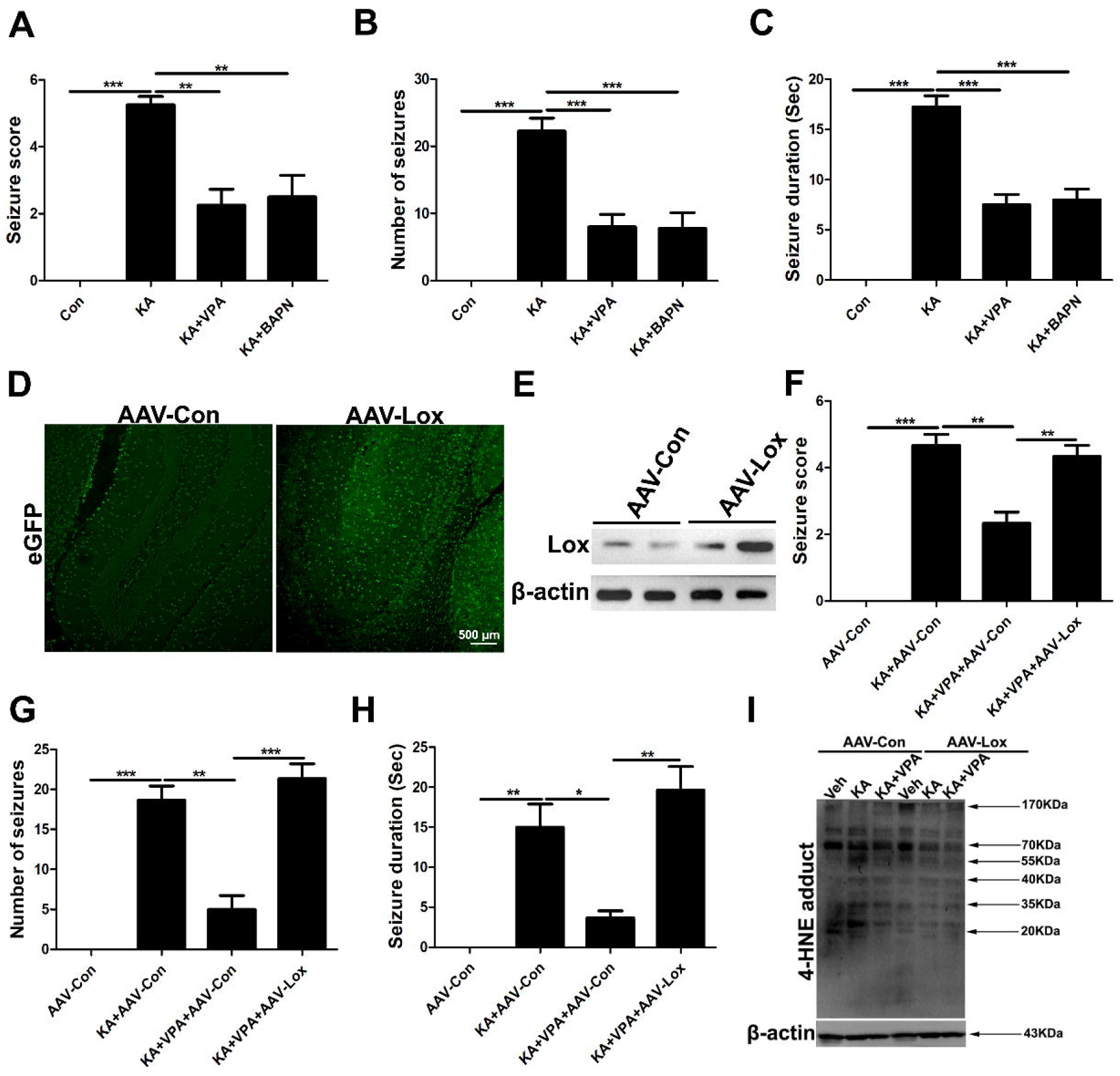
Overexpression of Lox reversed the neuroprotective of VPA in kainic acid-induced epilepsy mice model. **(A-C)** VPA (250 mg/Kg) and BAPN (100 mg/Kg) could reduce the level of seizure score, number of seizures, and average seizure duration (Sec) in kainic acid (250 ng) induced epilepsy mice model, n=6. **(D)** Representative immunofluorescence images showed that the expression of eGFP was observed in hippocampus tissue of Lox overexpression group, scar bar=500 μm. **(E)** Western blot analysis showed that the expression of Lox in hippocampus tissue of pAAV-SYN-Lox-P2A-EGFP-3FLAG group was significantly higher than that in pAAV-SYN-EGFP-3FLAG group. **(F-H)** Pretreatment with VPA (250 mg/Kg) could not ameliorate the level of seizure score, number of seizures, and average seizure duration (Sec) caused by kainic acid (250 ng) in Lox overexpression group, n=6. **(I)** Pretreatment with VPA (250 mg/Kg) could not reduce the production of 4-HNE caused by kainic acid (250 ng) in Lox overexpression mice model, *p < 0.05, **p < 0.01 and ***p < 0.0001.

## 4. Discussion

In our present work, we found that ferroptosis was involved in both epilepsy animal models and glutamate-induced cell models. And the neuroprotective effect of VPA in the epilepsy model is related to the inhibition of ferroptosis. In addition, further mechanistic research indicates that VPA can play a neuroprotective role in KA-induced epilepsy model by inhibiting Lox-mediated ferroptosis.

In China, epilepsy is the second most common neurological disease, whose morbidity is second to cerebrovascular disease. As a kind of chronic disease, epilepsy, not only causes serious physical and psychological harm to the patients themselves, but also increases the social and economic burden. Therefore, it is of utmost importance to explore effective strategies in epilepsy. Drug therapy is a major way for curing epilepsy. At present, there are more than 20 kinds of antiepileptic drugs used in the treatment of epilepsy^[32]^. Different types of seizures require the use of different antiepileptic drugs: phenobarbital and phenytoin are often used as the first choice for treating epileptic seizures^[33]^. And for absence seizure, ethosuximide is often used as the primary therapeutic drug^[34]^. As a classic antiepileptic drug, VPA has a good therapeutic effect on many types of epilepsy. The antiepileptic mechanism of VPA is not clear. Previous studies have indicated that VPA may play an antiepileptic effect by inhibiting Na^+^ channel activity and enhancing the transmission of γ-aminobutyric acid^[35,36]^. Besides, more and more studies believe that VPA has certain neurotrophic and neuroprotective effects. Because of its safety and therapeutic effect, VPA is also widely used in the prevention of seizures after neurosurgical operations such as craniocerebral trauma. However, the specific neuroprotective mechanism in epilepsy is still unclear. Therefore, exploring the neuroprotective mechanism of VPA in epilepsy will provide new ideas for the treatment of epilepsy.

Previous studies have shown that VPA can play a neuroprotective role in central nervous system diseases through antioxidant effects. In chronic neurological diseases such as spinal cord injury, stroke and traumatic brain injury, VPA can protect neurons from oxidative damage by inhibiting the production of mitochondrial ROS, endoplasmic reticulum stress and lipid peroxidation^[13,37]^. In our research, we also found that VPA could inhibit lipid peroxidation in a mouse model of epilepsy, thereby exerting a neuroprotective effect in epilepsy. Recent studies have found that lipid peroxidation can mediate a new type of cell death-ferroptosis. A large number of studies have shown that ferroptosis is involved in the pathogenesis of various central nervous system diseases including epilepsy^[38]^. In the first part of our research, we found that VPA and ferroptosis inhibitors could significantly improve seizures in epileptic mice. This suggests that ferroptosis is involved in the KA-induced epilepsy model. The neuroprotective effect of VPA in this model may be related to the inhibition of ferroptosis. To further clarify the relationship between the neuroprotective effect of VPA and ferroptosis in this model. We tested ferroptosis biomarkers in the mouse hippocampus of epilepsy model. 4-HNE and PTGS2 are commonly used biomarkers in ferroptosis research. 4-HNE is the most representative aldehyde group product produced by ω6 polyunsaturated fatty acids such as arachidonic acid and linoleic acid through multiple lipid chain cleavage reactions in lipid peroxidation, which can reflect the occurrence of lipid peroxidation^[39]^. Lipid peroxidation-mediated cell damage is indispensable for the occurrence of ferroptosis, so the increase of 4-HNE can indicate the occurrence of ferroptosis to some extent. PTGS2 is a gene encoding cyclooxygenase-2, which is significantly upregulated during the occurrence of ferroptosis, and it is a downstream biomarker of ferroptosis^[40]^. Our results showed that, compared with the control group, 4-HNE and PTGS2 levels in the KA model group were significantly increased, while the VPA and ferroptosis inhibitor groups were significantly inhibited in the model group. This further indicates that ferroptosis is implicated in the KA-induced epilepsy model, and the protective effect of VPA in the KA-induced epilepsy model is related to the inhibition of ferroptosis. We further used glutamate-induced HT22 cell damage model to explore the specific molecular mechanism of VPA underlying the inhibition of ferroptosis. Glutamate is a major excitatory neurotransmitter in the nervous system. After epileptic seizures, glutamate in the brain begins to accumulate in large quantities, causing neuronal damage^[28]^. And the HT22 cell is a good model for studying oxidative damage caused by glutamate^[30]^. Moreover, there are some studies supporting that ferroptosis is involved in glutamate-induced HT22 cell damage^[31]^. Our previous study also indicated the occurrence of ferroptosis in the glutamate-induced HT22 cell damage. Based on the above considerations, this study selected glutamate-induced HT22 cell damage model for in vitro molecular mechanism of VPA neuroprotection. We found that pretreatment of VPA and ferroptosis inhibitors could play a significant protective role against glutamate-induced cell damage. For further verifying the above findings, we tested the ferroptosis-related indicators after VPA pretreatment in a glutamate-induced cell damage model. Finally, we found that: VPA could significantly reduce the production of lipoid ROS, 4-HNE and MDA. In addition, in the HT22 cell damage model induced by the ferroptosis-specific inducer elastin, we also found that VPA exerted significant neuroprotection and reduced the level of ferroptosis-related biomarkers. It suggests that the protective effect of VPA on glutamate-induced HT22 cell damage is closely related to the inhibition of ferroptosis. In the previous research, our research group found that Lox is a key regulator of the occurrence of ferroptosis in epilepsy. To investigate whether Lox is a potential target for VPA to inhibit ferroptosis, we tested Lox mRNA and protein levels in a glutamate-induced HT22 cell damage model. We found that: consistent with the previous upregulation of Lox expression in the KA model, Lox expression also increased significantly in the glutamate-induced HT22 cell injury model, and VPA pretreatment could significantly reduce its expression level. At the same time, our results also found that: BAPN, specific inhibitor of Lox, could protect HT22 cells from glutamate-induced damage and inhibit the increase of ferroptosis indicators. These results indicate that Lox may be the key target for VPA to inhibit ferroptosis. In this study, an adenoviral overexpression vector was used to construct a Lox overexpression mouse model to further explore in vivo whether VPA plays a neuroprotective role in thalamic acid-induced epilepsy models by inhibiting Lox. We found that, in Lox overexpressed mice, VPA did not inhibit the KA-induced seizures in mice. Although previous reports have pointed out that ferroptosis is involved in the development of neurodegenerative diseases such as Parkinson’s disease and Alzheimer’s disease^[38]^, little is known about the role of ferroptosis in the process of epilepsy. This study revealed that ferroptosis was involved in seizures and played a key role in the neuroprotection of VPA. Besides, our research also shows that VPA can play a neuroprotective mechanism in epilepsy by inhibiting Lox-mediated ferroptosis. However, the specific molecular mechanism of the regulation of Lox expression by VPA in KA-induced epilepsy model, the function of Lox in ferroptosis, and the molecular biological process require further research in the future. Manipulation of ferroptosis can be used as a promising therapeutic target for epilepsy, although the contribution of ferroptosis in the pathogenesis of various neurological diseases remains to be explored.

## 5. Conclusion

VPA plays a neuroprotective role in KA-induced epilepsy, which is related to the inhibition of ferroptosis. In vitro cell models confirm that Lox-mediated ferroptosis is involved in the protective effect of VPA on glutamate-induced cell damage models. Our findings showing that Lox overexpression inhibits the neuroprotection of VPA against epileptic seizures suggest that VPA plays a neuroprotective role in the -induced epileptic seizures by inhibiting Lox-mediated ferroptosis.

## Conflict of interest

The authors declare that they have no potential conflict of interest.

## Acknowledgments

This work is partially financially supported by the National Natural Science Foundation of China (Nos. 81671293 and 81974502).

## Notes

### Competing Interest Statement

The authors have declared no competing interest.

